# Anti-HIV-1 HSPC-based gene therapy with safety kill switch to defend against and attack HIV-1 infection

**DOI:** 10.1101/2024.11.13.623476

**Authors:** Qi Guo, Keval Parikh, Jian Zhang, Alexander Brinkley, Grace Chen, Natnicha Jakramonpreeya, Anjie Zhen, Dong Sung An

## Abstract

Hematopoietic stem/progenitor cell (HSPC)-based anti-HIV-1 gene therapy holds promise to provide life-long remission following a single treatment. Here we report a multi-pronged anti-HIV-1 HSPC-based gene therapy designed to defend against and attack HIV-1 infection. We developed a lentiviral vector capable of co-expressing three anti-HIV-1 genes. Two are designed to prevent infection, including a short-hairpin RNA (CCR5sh1005) to knock down HIV-1 co-receptor CCR5 and a membrane anchored HIV-1 fusion inhibitor (C46). The third gene is a CD4-based chimeric antigen receptor (CAR) designed to attack HIV-1 infected cells. Our vector also includes a non-signaling truncated human epidermal growth factor receptor (huEGFRt) which acts as a negative selection-based safety kill switch against transduced cells. Anti-HIV-1 vector-transduced human CD34+ HSPC efficiently reconstituted multi-lineage human hematopoietic cells in humanized bone marrow/liver/thymus (huBLT) mice. HIV-1 viral load was significantly reduced (1-log fold reduction, p <0.001) in transplanted huBLT mice. Anti-huEGFR monoclonal antibody Cetuximab (CTX) administration significantly reduced huEGFRt+ vector-modified cells (>4-fold reduction, p <0.01) in huBLT mice. These results demonstrate that our strategy is highly effective for HIV-1 inhibition, and that CTX-mediated negative selection can deplete anti-HIV-1 vector-modified cells in the event of unwanted adverse effects in huBLT mice.

## Introduction

Forty years after its discovery, HIV-1 infection remains a significant public health issue with a total of over 39 million cases and more than 1.3 million new cases globally in 2023 alone.^1^ Although antiretroviral therapy (ART) has significantly improved life expectancy and health of those living with HIV, it cannot cure HIV-1 infection.^2–6^ Life-long treatment is necessary due to the persistence of HIV-1 infection by producing latent HIV-1 viral reservoirs.^4–7^ Other shortcomings of ART include patient adherence, administrative availability, drug cost, and adverse side effects. Moreover, ART reduces but does not prevent all of the known complications of HIV-1 infection.^8–12^ A novel therapeutic approach for life-long remission without ART or elimination is critical to address these issues.^5,13^

Thus far, HIV-1 cure has only been achieved in a few patients who have undergone hematopoietic stem cell transplantation (HSCT) from HLA type fully or partially matched allogeneic CCR5Δ32/Δ32 homozygous donors.^14–19^ In these few patients, allogeneic transplantation with CCR5Δ32/Δ32 HSPC to treat underlying leukemia has also led to long-term HIV-1 remission without the need for ART after successful repopulation of immune cells lacking the HIV-1 co-receptor CCR5. These handful HIV-1 cure cases offer great hope for the development of an HSPC-based anti-HIV-1 gene therapy that provides long-term remission or cure for HIV-1 infection. However, CCR5Δ32/Δ32 homozygous mutation is found in less than 1% of the global population.^20,21^ Furthermore, allogeneic stem cell transplantation also requires HLA-matching,^16–18^ making it extremely difficult to identify a HLA type matched donor with CCR5Δ32/Δ32 homozygous mutation. Despite this, HSPC-based gene therapy to genetically modify autologous cells with anti-HIV-1 genes holds great promise to provide life-long remission or cure following a single treatment.^22–24^ Unlike HSCT, autologous HSPC-based gene therapy uses a patient’s own cells and hence does not require HLA-matching.^23,25^

Anti-HIV-1 HSPC-based gene therapy may require a multi-target approach to effectively inhibit HIV-1, similar to the combinatorial drug treatment strategy used in ART.^26,27^ We previously identified and proved the potent antiviral activity of a non-toxic short hairpin RNA against CCR5 (CCR5sh1005) to down regulate CCR5 expression by RNA interference to protect cells from HIV-1 entry.^28,29^ Although our CCR5sh1005 was efficient for down regulating CCR5 in human CD4+ T cells and HIV-1 inhibition through HSPC gene-modification in humanized BLT mice and rhesus macaques, it does not fully ablate CCR5 expression.^30–33^ We therefore added a membrane anchored anti-HIV-1 fusion inhibitor C46, which targets gp41 on HIV-1 virions to prevent fusion into host cells, a critical viral entry step.^30,31,33^ These anti-HIV-1 genes work synergistically to protect cells by inhibiting HIV-1 binding and fusion before viral integration to prevent the establishment of chronic HIV-1 infection. We previously demonstrated that dual anti-HIV-1 combinations (CCR5sh1005 and C46) improved HIV-1 inhibition compared to CCR5sh1005 alone and inhibited both R5-tropic and X4-tropic HIV-1.^33^

In addition to defending HSPC against HIV-1 infection by use of the aforementioned anti-HIV-1 genes, we reasoned that the potential for chronic remission of HIV-1 infection could be increased by simultaneously engineering a host immunological attack on HIV-1 infected cells. We incorporated a CD4-based anti-HIV-1 CAR that has demonstrated robust HIV-1 viral load reduction when expressed in HSPC, producing anti-HIV-1 CAR T cells to attack and eliminate HIV-1 infected cells.^34^ CD4-based anti-HIV-1 CARs are designed to bind a HIV-1 GP120 envelope glycoprotein on cell surface with the extracellular CD4 D1D2 HIV-1-binding domain and transmit signals through the intracellular CD3-ζ signaling domain to kill HIV-1 infected cells by T cell mediated cytotoxicities.^35,36^ D1D2CAR 4-1BB is a truncated version of the previously used CD4CAR and includes a 4-1BB costimulatory domain shown to enhance CAR-T cell function and proliferation compared to other anti-HIV-1 CAR-T cell variants *in vivo*.^37^ D1D2CAR 4-1BB also does not mediate HIV-1 infection and when coupled with anti HIV-genes such as CCR5sh1005 provide gene-modified cells with extra protection.^37^ CD4-based CARs has been co-expressed in dual combination lentiviral vectors with C46 or CCR5sh1005.^34,37^ We hypothesize that triple-expression of CCR5sh1005, C46, and D1D2CAR 4-1BB in HSPC will durably protect infection-susceptible progeny cells and target HIV-1 infected cells, thereby inhibiting 3 different steps in HIV-1 infection.

Despite a superior safety profile in our humanized mice and NHP studies,^34,37,38^ incorporating a negatively-selectable safety kill switch into our gene therapy could prove important for our approach. Anti-HIV-1 CAR T vector-modified cells may potentially cause unexpected health issues such as cytokine release syndrome and CAR-T cell related encephalopathy syndrome in hosts, as seen in cancer immunotherapy.^39–41^ To improve safety of our HSPC-based anti-HIV-1 gene therapy, we incorporated a safety kill-switch by co-expressing the non-functional truncated form of human epidermal growth receptor (huEGFRt), which can be targeted with the clinically available chimeric immunoglobulin G1 anti-EGFR monoclonal antibody (mAb), Cetuximab (CTX) (Erbitux™).^42,43^ Administration of CTX can negatively select huEGFRt expressing vector-modified cells *in vivo* through antibody-dependent cellular cytotoxicity (ADCC) and complement dependent cytotoxicity (CDC). To prevent off-target activation of CAR T-cells while retaining a marker function for the tracking of vector-modified cells via mAb staining and flow cytometry analysis, huEGFRt lacks the extracellular ligand binding domains I and II and the entire cytoplasmic tail necessary for signaling in EGFR.^42^

In this report, we developed a multi-pronged anti-HIV-1 lentiviral vector with a safety kill switch for efficient HSPC transduction and expression of factors capable of protecting cells against and attacking HIV-1 infection. We investigated the ability of our newly developed anti-HIV-1 gene lentiviral vectors to genetically modify human CD34+ HSPC, and the ability to transplant and engraft these vector-modified cells to inhibit HIV-1 infection *in vivo* in huBLT mice. Furthermore, we investigated a safety kill switch by CTX-mediated negative selection of huEGFRt+ vector-modified cells. Together, these elements work together to provide a robust and safe anti-HIV-1 HSPC-based gene therapy strategy.

## Results

### Development of lentiviral vectors with a safety kill switch to defend and attack HIV-1 infection

We developed two new lentiviral vectors (M1 vector: MNDU-anti-HIV-1-huEGFRt and U1 vector: UbC-anti-HIV-1-huEGFRt) to effectively co-express three anti-HIV-1 genes to defend against and attack HIV-1 infection and to express the huEGFRt cell surface marker as a safety kill switch (Figure 1A). We expressed CCR5sh1005 from a transcriptionally weaker H1 RNA Polymerase III promoter to avoid toxic effects of shRNA overexpression, as previously described.^28,29^ We first examined a modified Moloney Murine Leukemia virus long terminal repeat promoter (MNDU) and a Ubiquitin C (Ubc) RNA polymerase II promoter to optimize the co-expression of D1D2CAR 4-1BB and huEGFRt expression in M1 vector and U1 vector, respectively. These two transgenes were linked by a self-cleaving T2A sequence for equimolar expression. C46 was expressed from a shorter version of the eukaryotic translation elongation factor 1α (EF1alpha) promoter to maintain efficient expression as the last transgene inserted near the 3’LTR. Despite the multiple promoters and transgenes, the titers of our newly developed anti-HIV-1 vectors in 293T cells were high (2.75×10^8^ ± 3.03×10^7^ IU/mL for the M1 vector and 1.00×10^8^ ± 1.95×10^7^ IU/mL for the U1 vector) (Figure 1B), which is consistent with our previously developed lentiviral vectors.^31^ Normalized % CCR5 expression was reduced to 76.1% and 69.8% in M1 and U1 vector transduced huEGFRt+ MT4-CCR5 cells, respectively, compared to the normalized %CCR5 in Non-CCR5sh1005 vector transduced huEGFRt+ cells (100%) (Figure 1C). Mean fluorescent intensity (MFI) of CCR5 expression in huEGFRt+ cells was reduced to 1491 and 1238 in M1 and U1 vector transduced huEGFRt+ cells, respectively, compared to 10526 in Non-CCR5sh1005 vector transduced huEGFRt+ cells. These results show CCR5 expression was efficiently down-regulated in M1 and U1 vector-transduced huEGFRt+ MT4-CCR5 cells. We noticed that MFI of huEGFRt expression were lower in M1 (1197) and U1 (1139) vector transduced cells than that of non-CCR5sh1005 vector transduced cells (5665) (Fig 1C), suggesting that huEGFRt expression might be compromised due to 4 multiple transgene expressions from one vector. C46 was efficiently expressed in both M1 (94.6% C46+) and U1(98.9% C46+) vector-transduced MT4-CCR5 cells (Figure 1D). D1D2CAR 4-1BB and huEGFRt were efficiently co-expressed in vector-transduced human primary CD8+ T cells by both the M1 vector (36.4%) and U1 vector (49.1%) (Figure 1E). These results show that our newly developed vectors are highly efficient for co-expressing three anti-HIV-1 genes and huEGFRt in human T-cell line and primary human T cells *in vitro*. Both R5-tropic HIV-1_NFNSX- SL9_ and X4-tropic HIV-1_NL4-3_ infection were inhibited in M1 and U1 vector transduced MT4- CCR5 cell line *in vitro* (Figure 1F). To examine cell killing activity of M1 and U1 vector-transduced CD8+ CAR T cells, we performed cytotoxic T lymphocyte (CTL) assays by co-incubating vector-transduced CD8+ T cells with either ACH2 cells stimulated to express high levels of HIV-1 envelope (Env+) or unstimulated ACH2 cells (Env-). We observed approximately 60% specific killing for both M1 and U1 vector-transduced CAR T cells at an E:T ratio of 5:1 (p <0.01, p <0.05, respectively) (Figure 1G). These results reinforce the potential of D1D2CAR 4-1BB to specifically target and induce cellular cytotoxicity in HIV-1 envelope expressing cells. Altogether, these results demonstrate successful construction of a multi-pronged anti-HIV-1 lentiviral vector that can block both R5- and X4-tropic HIV-1 infection *in vitro* and direct a cellular immune response against infected cells via a chimeric antigen receptor.

**Figure 1.**
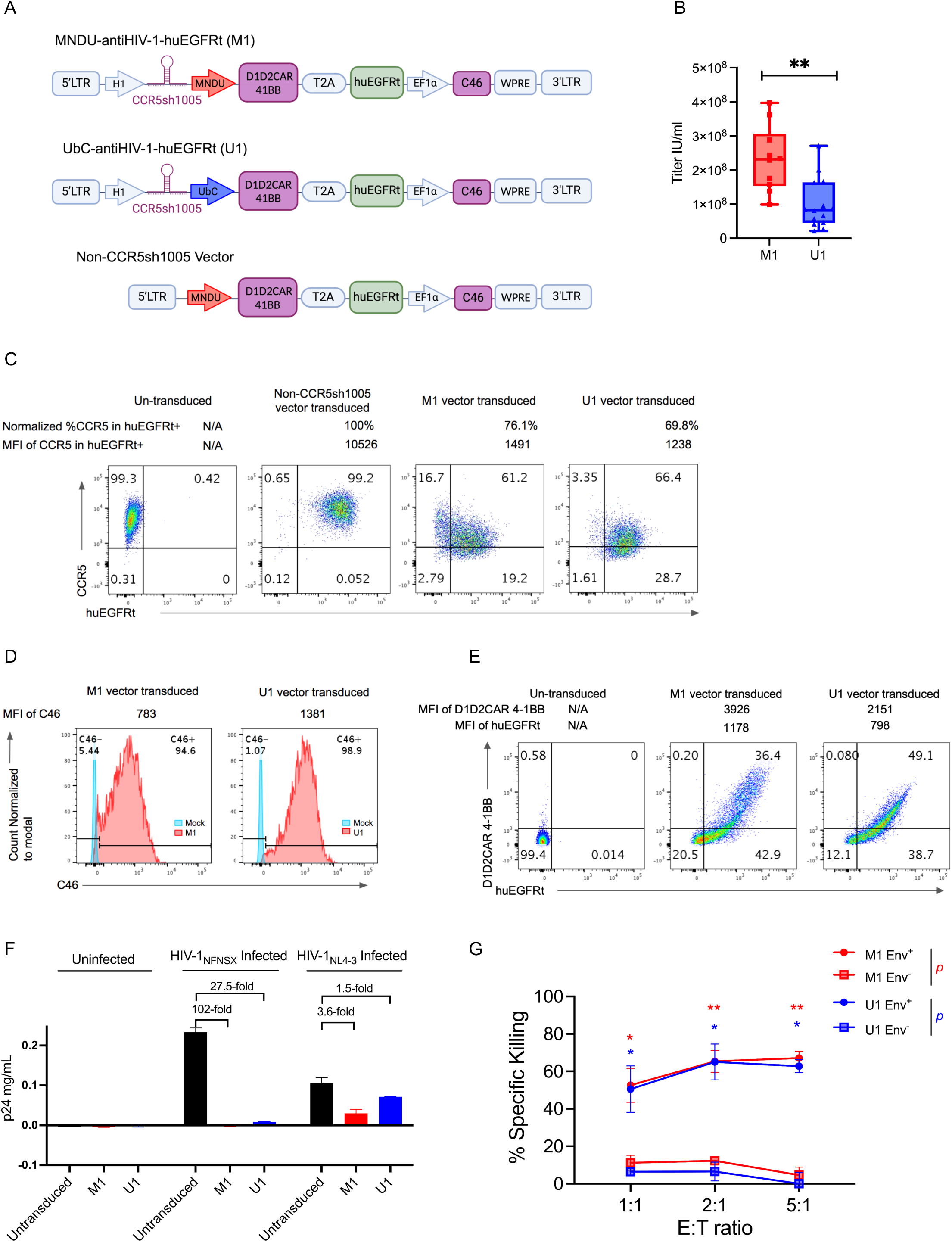
Development of a multi-pronged anti-HIV-1 lentiviral vector with safety kill switch to defend against and attack HIV-1 infection. (A) Design of novel lentiviral vectors expressing triple anti-HIV-1 genes (CCR5sh1005, D1D2CAR 4-1BB, C46) and a selectable huEGFRt gene. The M1 vector uses an MNDU promoter and the U1 vector uses a Ubc promoter for D1D2CAR 4-1BB and huEGFRt expression, respectively. These vectors also includes ΔLTR: self-inactivating, U3 enhancer and promoter deleted long terminal repeat, H1: H1 RNA polymerase III promoter, MNDU: murine leukemia virus (MuLV) long terminal repeat promoter, UbC: Ubiquitin C RNA polymerase II promoter, T2A: 2A self-cleaving peptide, huEGFRt: truncated nun-functional human epidermal growth factor receptor, EF1α: human elongation factor 1 alpha promoter, WPRE: woodchuck hepatitis virus posttranscriptional regulatory element. Non-CCR5sh1005 vector includes ΔLTR, MNDU, D1D2CAR 4-1BB, T2A, huEGFRt, C46 and WPRE. (B) Vector titer measured by infectious units/milliliter (IU/ml) determined based on percent huEGFRt transgene expression in 293T cell line stained with CTX-PE mAb and measured by flow cytometry at day 3 post-transduction. (C) CCR5 downregulation by CCR5sh1005 in MT4-CCR5 cell lines. MT4-CCR5 cells were transduced at MOI 1 with vectors M1 and U1. Untransduced and non-CCR5sh1005 vector transduced MT4-CCR5 cells were used as a negative control. 4 days post-vector transduction, CCR5 and huEGFRt were stained with mAbs and measured by flow cytometry. Normalized CCR5 expression within huEGFRt positive population was calculated based on the following formula: ([%CCR5+/huEGFRt+population]/([%CCR5+/huEGFRt+population] + [%CCR5-/huEGFRt+population])×100%) and mean fluorescent intensity of CCR5 expression in huEGFRt positive population (MFI) is indicated on the top of representative flow plot. (D) C46 cell surface expression and MFI in MT4-CCR5 cell line at 4 days post-transduction. MT4-CCR5 cells were transduced at MOI 1 with vectors M1 and U1. Mock is untransduced negative control cells. C46 was stained with mAb and measured by flow cytometry. (E) HuEGFRt and D1D2CAR 4-1BB surface protein expression in primary human CD8+ T cells 4 days post-transduction. CD8+ T cells isolated from healthy-donor PBMCs were transduced with M1 and U1 vectors, respectively (MOI 10). Untransduced primary human CD8+ T cells were used as a negative control. HuEGFRt and D1D2CAR 4-1BB expression was stained with mAb and measured by flow cytometry. (F) *In-vitro* HIV-1 inhibition by M1 and U1 vector-transduced MT4-CCR5 cells. Untransduced MT4-CCR5 cells were used as a negative control. MT4-CCR5 cells were challenged with R5 tropic HIV-1_NFNSX_ (MOI 1) or X4 tropic HIV-1_NL4-3_ (MOI 0.005) virus. p24 capsid protein levels in cell culture supernatant were determined by p24 ELISA assay 7 days post HIV-1 challenge and used to assess inhibition ability. Data shows results from two independent experiments. (G) *In-vitro* specific killing of HIV-1 envelope expressing ACH2 cells. PMA/ionomycin stimulated (Env+) or unstimulated (Env-) ACH2 cells were co-cultured with vector-transduced human primary CD8+ T cells at 1:1, 2:1, and 5:1 Effector:Target cell (E:T) ratio overnight. %specific killing was calculated by (%live gag+ ACH2 cells with untransduced CD8+ cell—%live gag+ ACH2 cells with vector transduced CD8+ cells)/ %live gag+ ACH2 cells with untransduced CD8+ cell then normalized by %gag+ in ACH2 cells. Data shows Mean ± SEM from a single experiment performed in triplicates. Mann-Whitney U test was performed to calculate significance. *p <0.05 and **p <0.01.

## Efficient engraftment of vector-modified HSPC for HIV-1 viral load reduction in huBLT mice

We next investigated the efficiency of vector transduction and transplantation of human CD34+ HSPC to assess the engraftment, multi-lineage hematopoietic cell reconstitution, transgene expression and HIV-1 inhibition *in vivo* in the huBLT mouse model (Figure 2A). M1 and U1 vectors efficiently transduced human fetal liver derived CD34+ HSPC (FL-CD34+ HSPC) at multiplicity of infection (MOI) 3 *ex vivo*. D1D2CAR 4-1BB+/huEGFRt+ co-expressing cell population reached 79.9% and 51.7% in M1 and in U1 vector-transduced FL- CD34+ HSPC at day 4 post vector transduction (Figure 2B). The MFI of D1D2CAR 4-1BB (13080) and huEGFRt (14806) expression in the M1 vector-transduced HSPC was notably higher than that of the U1 vector-transduced HSPC D1D2CAR 4-1BB (889) and huEGFRt (1344), similar to the vector-transduced human primary CD8+ T cells (Figure 1E). Multi-lineage colony formation in *ex vivo* culture showed similar % of multi colony-forming units between untransduced, M1, and U1 vector-transduced FL-CD34+ HSPC *ex vivo* (Figure S1). After transplantation of vector-transduced FL-CD34+ HSPC in huBLT mice, total CD45+, CD3+, CD4+ and CD8+ T, and CD19+ B multilineage human hematopoietic cells were reconstituted and continued to expand in peripheral blood in huBLT mice, as previously reported.^44,45^ There were no significant differences between untransduced control, M1, and U1 vector-transduced FL-CD34+ HSPC transplanted huBLT mouse groups (herein referred to as untransduced, M1, U1 huBLT mice, respectively) (Figure 2C). The average vector DNA copies/human cell were higher in M1 vs U1 huBLT mice (∼2 copies/cell vs ∼1 copy/cell) and stably maintained during the experiment (Figure 2D). The higher vector copy number in M1 huBLT mice compared to U1 huBLT mice could be attributed to a higher packaging efficiency of a lentiviral vector with an MNDU promoter, as previously reported.^46^ In a subsequent experiment, similarly high vector copy levels (∼1.5 copies/cell) were detected in M1 huBLT mice (Figure S2). Absolute numbers of huEGFRt+ human CD3+, CD4+ and CD8+ T cells, but not CD19+ B cells significantly increased in M1 vector-modified huBLT mice compared to U1 vector-modified and untransduced huBLT mice after 4 to 6-weeks post-R5 tropic HIV-1_NFNSX-SL9_ challenge in peripheral blood (Figure 2E), suggesting CCR5sh1005 and C46 may provide protection for M1 vector-modified human CD4+ T cells for selective growth advantage, and D1D2CAR 4-1BB may provide a CAR-dependent proliferation advantage. Percentage of huEGFRt+ cells did not show increase in peripheral blood in M1 huBLT mice and in U1 huBLT mice because huEGFRt- cells also increased in the huBLT mice (Figure S3, S4, and S5) as total human hematopoietic cells continue to increase in huBLT mice (Figure 2C).

**Figure 2.**
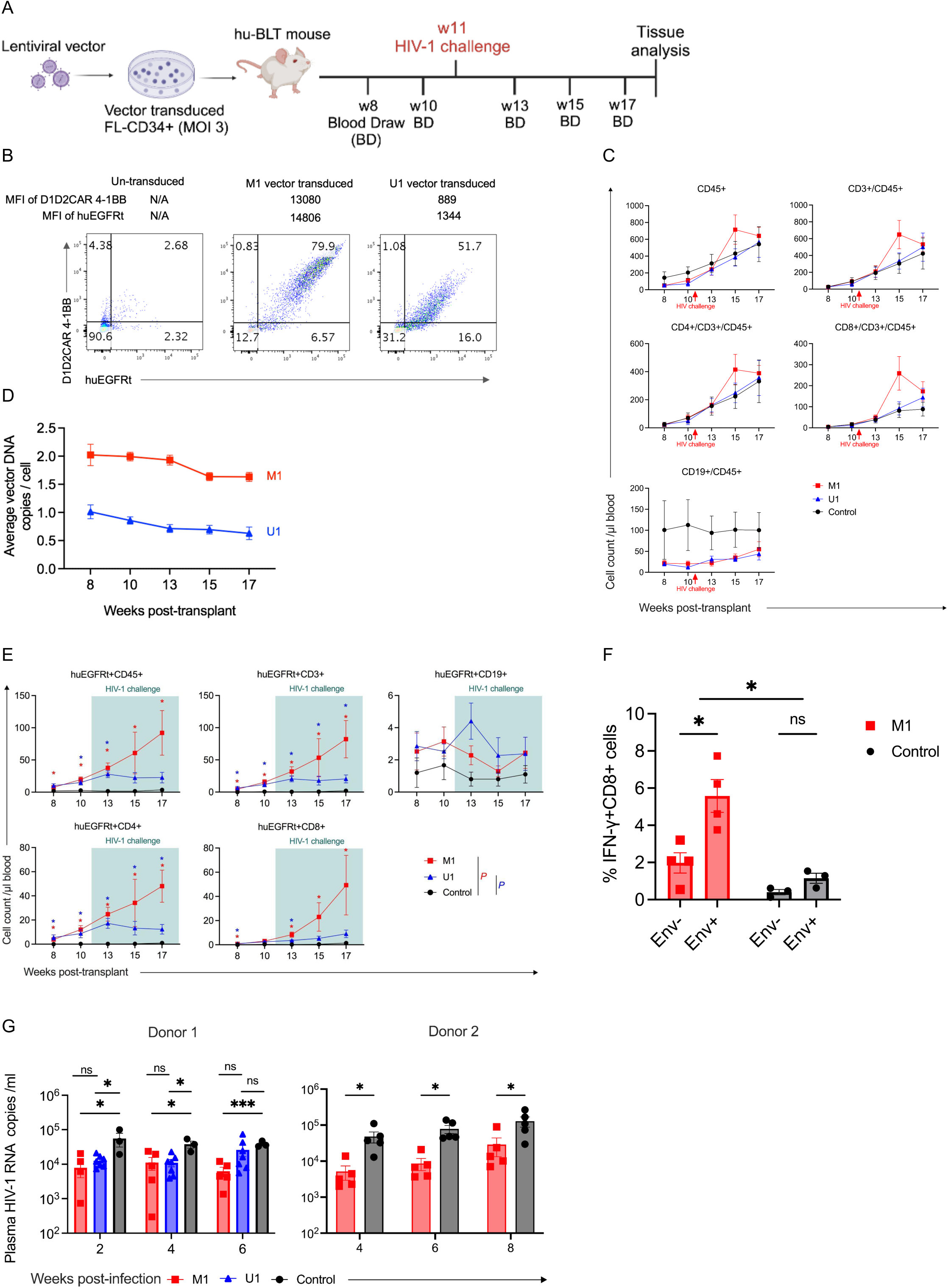
Efficient human HSPC vector-modification, transplantation and multi-lineage human hematopoietic cell reconstitution in huBLT mice. (A) Experimental design for the investigation of M1 and U1 vectors in NSG huBLT mice. Human FL-CD34+ cells were transduced with M1 or U1 vectors at MOI 3 on Day -1. NSG huBLT mice were conditioned with 270 cGy of sub-lethal body irradiation from a Cesium-137 source. Mice were transplanted with the vector transduced FL-CD34+ HSPC and human thymus tissue on Day 0. Mice were challenged with R5-tropic HIV-1_NFNSXSL9_ (200ng p24/ mouse) at 11 weeks post-transplant. (B) HuEGFRt and D1D2CAR 4-1BB transgene expression in vector transduced FL-CD34+ cells in *ex-vivo* culture. HuEGFRt and D1D2CAR 4-1BB co-expressing population was determined by mAb staining and flow cytometry 4 days post-vector transduction. (C) Human multilineage hematopoietic cell reconstitution in peripheral blood from 8 weeks post-vector transduced HSPC transplant. Cell surface markers of human lymphocytes (CD45), T-cells (CD3, CD4, and CD8) and B-cells (CD19) were determined by mAb staining and flow cytometry. Dots and error bars show Mean ± S.E.M, respectively. (D) Vector-marking levels were determined in peripheral blood cells from 8 weeks to 17 weeks post-transplant by digital PCR. Average vector DNA copies were calculated by VCN = (WPRE DNA copies in vector DNA/ul)/(human β-globin copies/ul/2). Dots and error bars show Mean ± S.E.M, respectively. (E) Human multilineage hematopoietic cell expansion within huEGFRt expressing population after HIV-1 infection. Expression of huEGFRt was determined by mAb staining and flow cytometry. Cell surface markers of human lymphocytes (CD45), T-cells (CD3, CD4, and CD8) and B-cells (CD19) were also determined by mAb staining and flow cytometry and gated within the huEGFRt positive population. Mice were challenged with HIV-1 at 11 weeks-post transplant (not noted in this figure). Dots and error bars show Mean ± S.E.M, respectively. Mann-Whitney U test was performed to calculate significance, *p <0.05. (F) E*x vivo* cytokine production measured by cytokine release assay. CD8+ T splenocytes from M1 huBLT mice were co-cultured with Env+ target cells (PMA/ionomycin activated ACH2 cells) or unstimulated Env- cells (medium only) as a negative control *ex vivo*. Data was collected from our replicate huBLT mice experiment (donor 2). Cells were collected at time of mouse sacrifice at week 20 post-transplant. Cytokine expression was measured by flow cytometry. Dots and error bars show Mean ± S.E.M, respectively. *t*-test with Holm-Šídák adjustment was performed to calculate significance. *p <0.05. (G) Viral loads were measured as HIV-1 RNA copies per mL in mouse plasma every 2 weeks post- HIV-1 challenge by digital PCR in 2 different sets of experiments using 2 human CD34+ HSPC donors (Donor 1 and Donor 2). HuBLT mice groups were transplanted with either M1- (n = 5 in both experiments) or U1-transduced (n = 7) HSPC. Untransduced huBLT mice were used as a negative control in both experiments (n = 3 in first experiment, and n = 5 in replicate experiment). Data were shown in Mean ± S.E.M. *t*-test with Holm-Šídák adjustment was performed to calculate significance. NS = not significant, *p <0.05, and ***p <0.001.

To examine if vector-modified cells from the huBLT mice could respond to HIV-1 envelope protein, we performed *ex vivo* cytokine release assays using HIV-1 Env+ ACH2 cells as targets. When mixed with HIV-1 Env+ ACH2 cells, human CD8+ splenocytes from M1 huBLT mice exhibited significantly higher IFN-γ expression compared to CD8+ splenocytes from M1 huBLT mice mixed with HIV-1 Env- ACH2 cells (∼3 fold increase, p <0.05) and CD8+ splenocytes from control untransduced huBLT mice mixed with HIV-1 Env+ ACH2 cells (∼5 fold increase, p <0.05) consistent with an HIV-1 envelope-specific cytokine response (Figure 2F, Figure S6). The HIV-1 plasma viral load was significantly reduced (p <0.001, 1-log reduction) for 6 weeks post HIV-1 challenge in M1 huBLT mice compared to untransduced huBLT mice, which served as our negative control. We also observed reduction of HIV-1 viral load for up to 4 weeks post HIV-1 challenge in U1 huBLT mice (∼4 fold reduction, p <0.05), but viral load reduction was not significant at 6 weeks post infection. Because the M1 vector showed more significant viral load reduction in our donor 1 experiment, we further investigated the M1 vector and validated the effectiveness of HIV-1 viral load reduction in M1 huBLT mice in a repeat experiment with donor 2 (∼1 log-fold reduction, p <0.05) (Figure 2G). These results demonstrate that our multi-pronged anti-HIV-1 HSPC-based gene therapy strategy with M1 vector can achieve efficient *ex vivo* CD34+ cell transduction, support multi-lineage human hematopoietic cell reconstitution, stable transgene expression and greater viral load reduction compared to the U1 vector in huBLT mice.

### CTX-mediated negative selection of huEGFRt+ vector-modified cells as a safety kill switch

Although adverse effects have not been reported in anti-HIV-1 HSPC-based gene therapy preclinical studies or in clinical trials, potential adverse side-effects from lentiviral vector transduced HSPC or the induction of anti-HIV-1 CAR T cells must be approached prospectively. We therefore incorporated a safety kill switch into our anti-HIV-1 gene lentiviral vector, huEGFRt, triggered by the cognate CTX antibody. We investigated CTX-mediated negative selection of huEGFRt+ vector-modified cells in huBLT mice (Figure 3A). In our first experiment, we observed transient reduction of huEGFRt+ vector-modified cells in M1 huBLT mice (Figure S7). Since reconstituted human immune function in humanized mouse models is suboptimal, we hypothesized that our initial modest results were due to the limited number of human NK cells in huBLT mice.^47,48^ To enhance the number of functional human NK cells for ADCC, we injected human NK cells and an IL-15 expressing lentiviral vector to promote survival and function of human NK cells. In this augmented humanized mouse model, the percentage and absolute cell number of huEGFRt+ M1 vector-modified cells were significantly reduced following CTX treatment. We observed substantial reductions in CD45+, CD3+, CD4+, CD8+, and CD4+/CD8+ multi-lineage hematopoietic cells in peripheral blood of CTX-treated animals compared to CTX-untreated mice after 1 week of CTX injections; this difference persisted for 4 weeks (∼13 fold reduction, p <0.01, and ∼13 fold reduction, p <0.05, respectively, averaged across all cell lineages at week 4 post-CTX treatment) (Figure 3B and 3C). HuBLT mice were euthanized at 4 weeks post CTX injections; huEGFRt+ vector-modified cells were significantly reduced in spleen and BM (∼ 9 fold reduction, p <0.01, and ∼ 2.5 fold reduction, p <0.01, respectively, averaged across all cell lineages at week 4 post-CTX treatment) in CTX-treated vs CTX-untreated control M1 huBLT mice (Figure 3D, Figure S8). Within HSPC population, huEGFRt+ vector-modified CD34+/CD90+/CD38- HSPC were likewise significantly reduced (∼3 fold reduction p <0.05) in the BM of CTX-treated M1 huBLT mice (2.12% ± 1.20%) compared to CTX-untreated M1 huBLT mice (21.67% ± 7.92%) (Figure 3E and 3F). HuBLT mice remained healthy in CTX-treated and untreated groups, suggesting no apparent health adverse effects (Figure S9).

**Figure 3.**
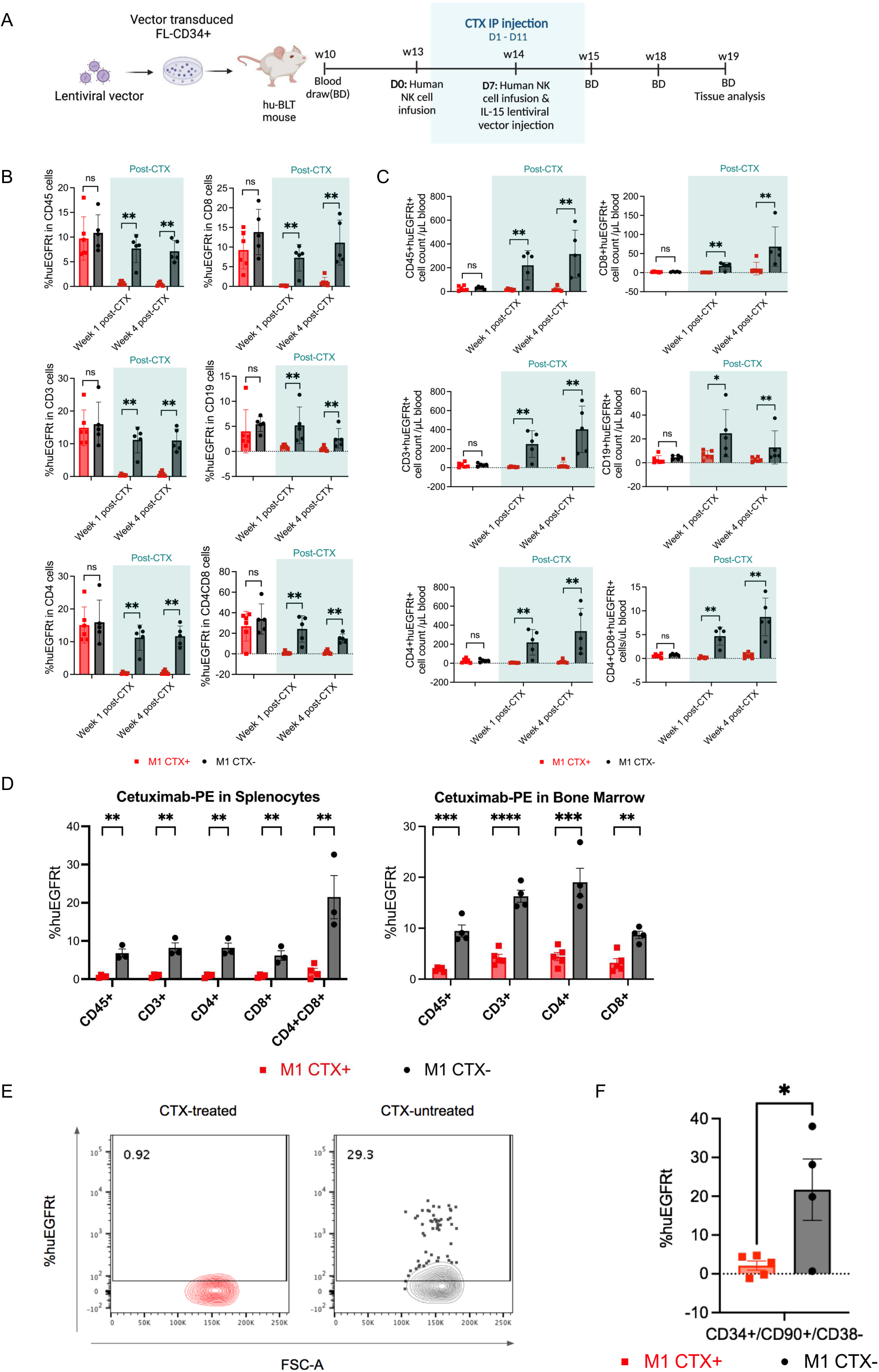
Cetuximab-mediated negative selection of huEGFRt expressing vector-modified human hematopoietic cells in huBLT mice. (A) Experimental design for the investigation of CTX-mediated negative selection of huEGFRt+ vector-modified human cells. CTX treatment group M1 huBLT mice (n=5) were injected with 1mg per mouse intraperitoneally for 11 consecutive days. 5x10^6^ human natural killer (NK) cells per mouse were injected retro-orbitally (RO) one day before first treatment (D0) and on D7 during CTX treatment. IL-15 expressing lentiviral vectors (2.5x10^6^ IU) were injected RO on day 7 of CTX treatment (2.5x10^5^ IU/mouse). CTX untreated group (n=4) served as a negative control. (B) HuEGFRt expression level in multilineage human peripheral blood cells (CD45+, CD3+, CD19+, CD4+, and CD8+) in CTX treated and untreated M1 huBLT mice. Blood was collected 1 week before CTX treatment, and 1 and 4 weeks post-onset of CTX treatment. Samples were stained with mAbs and measured by flow cytometry. HuEGFRt expression was measured by flow cytometry using mAb CTX-PE. Dots and error bars show Mean ± S.E.M, respectively. Mann-Whitney U test was performed to calculate significance. NS = not significant, and **p <0.01. (C) Absolute multilineage huEGFRt+ cell count in CTX-treated and untreated M1 huBLT mice in peripheral blood. Blood was collected at weeks 13, 15, and 18 post-transplantation (1 week pre-CTX treatment, and 1 and 4 weeks post-CTX treatment, respectively). HuEGFRt and surface markers of human lymphocytes (CD45), T-cells (CD3, CD4, and CD8) and B-cells (CD19), were stained by mAbs and measured by flow cytometry. Mann-Whitney U test was performed to calculate significance. NS = not significant, *p <0.05, **p <0.01, and ***p <0.001. (D) HuEGFRt expression level across multilineage human T-cell cell populations (CD45+, CD3+, CD4+, CD8+, and CD4+CD8+) in spleen and bone marrow tissue collected from CTX treated and untreated M1 huBLT mice. Samples were stained with mAb CTX-PE. Dots and error bars show Mean ± S.E.M, respectively. *t*-test with Holm-Šídák adjustment was performed to calculate significance. **p <0.01, ***p <0.001 and ****p <0.0001. (E) Representative flow cytometry data showing huEGFRt expression within CD34+/CD90+/CD38- population of bone marrow cells in CTX treated and untreated M1 huBLT mice. Samples were stained with mAb CTX-PE and measured by flow cytometry. (F) Cumulative data showing huEGFRt expression within CD34+/CD90+/CD38- population of bone marrow cells collected from CTX treated and untreated M1 huBLT mice. Samples were stained with mAb CTX-PE and measured by flow cytometry. % huEGFRt was normalized to background levels from mock transduced HSPC transplanted huBLT mice. Dots and error bars show Mean ± S.E.M, respectively. Student’s t-test was performed to calculate significance. *p <0.05.

Finally, we utilized Phycoerythrin (PE) conjugated CTX (CTX-PE) to stain huEGFRt+ cells and to analyze the level of expression by flow cytometry. To test whether CTX-mediated negative selection could impede detection of huEGFRt+ cells with the same antibody, we compared CTX-PE to another anti-huEGFR PE-conjugated mAb, Matuzumab (MTZ-PE), which binds to a different epitope on huEGFR (Figure S10).^49^ MTZ-PE staining confirmed that CTX- treated huEGFRt+ splenocytes from M1 huBLT mice were significantly reduced in multiple cell lineages (CD45+, CD3+, CD4+, CD8+) compared to CTX-untreated splenocytes from M1 huBLT mice (∼13-fold reduction, p <0.01 averaged across all cell lineages) (Figure S11). The lower %huEGFRt expression estimated by MTZ-PE staining may reflect the lower binding affinity of MTZ-PE than CTX-PE. Despite this difference, both MTZ-PE and CTX-PE stained CTX-treated M1 huBLT mice splenocytes showed significant reductions in huEGFRt expression. These results demonstrate that our CTX-mediated negative selection strategy is highly effective for depleting huEGFRt+ vector-modified human HSPC and progeny cells for diverse cell and gene therapies in huBLT mice.

## Discussion

In this study, we investigated a multi-pronged anti-HIV-1 HSPC based gene strategy to defend against and attack HIV-1 infection in humanized BLT mice. We developed a novel lentiviral vector that successfully co-expressed three anti-HIV-1 genes. These anti-HIV-1 genes include an shRNA against CCR5 HIV-1 co-receptor and C46 fusion inhibitor to protect cells against HIV-1 infection, and a truncated CD4-based CAR with 4-1BB costimulatory domain (D1D2CAR 4-1BB) to attack HIV-1 infected cells. We also incorporated huEGFRt, to allow for efficient negative selection of vector-modified cells as a safety kill switch in case of potential adverse effects. Our results demonstrate that vector-modified HSPC efficiently reconstituted anti-HIV-1 vector-modified cells and significantly reduced viral load *in vivo* in huBLT mice. We used huBLT mice since the development of human HSPC derived anti HIV-1 gene modified T cells occurs in the donor matched human thymus tissue. In other humanized mouse models, human T cell development occurs in mouse thymus and it is not efficient nor physiological due to the human leukocyte antigen (HLA) and mouse MHC mismatch.^44,50,51^ Administration of CTX, a clinically available anti-huEGFR monoclonal antibody, significantly reduced huEGFRt+ gene-modified cells, improving the safety of our anti-HIV-1 gene therapy strategy.

HSPC-based gene therapy has been investigated to achieve life-long remission or cure due to the potential of anti-HIV-1 gene modified HSPC to continuously provide HIV-1 protected immune cells.^22–24^ Unfortunately, the efficiency of gene modification in HSPC and the level of engraftment are not sufficient to achieve life-long remission with current technologies.^22–25^ If the engraftment and reconstitution is incomplete, remaining unprotected cells are subject to infection. CAR T-cells have emerged as a powerful immunotherapy for different forms of cancer.^52,53^ Anti-HIV-1 CAR gene can re-engineer host immune cells to target HIV-1 specific antigens such as gp120 on the surface of HIV-1 infected cells and elicit virus-specific cytotoxicity.^37,54,55^ This strategy subverts the necessity for complete engraftment of anti-HIV-1 vector modified HSPC, as anti-HIV-1 CAR T cells can attack HIV-1 infected cells. In addition to CCR5sh1005 and C46, we successfully developed a lentiviral vector capable of co-expressing a CD4-based D1D2CAR 4-1BB for efficient HSPC gene modification to achieve efficient viral load reduction.

We successfully incorporated a safety kill switch by co-expressing huEGFRt in our anti-HIV-1 HSPC based gene strategy to better prepare for potential adverse effects such as clonal outgrowth or malignant transformation of lentiviral vector transduced HSPC by random vector insertional mutagenesis, CAR T cell mediated cytokine release syndrome, or encephalopathy syndrome.^34,38,40,56–58^ We chose huEGFRt cell surface marker gene due to its relatively short cDNA sequence, allowing for its inclusion and efficient co-expression in a complex multi-pronged lentiviral vector. HuEGRt can be targeted with the clinically available chimeric immunoglobulin G1 anti-EGFR monoclonal antibody (mAb), Cetuximab (CTX) (Erbitux™) for efficient negative selection.^42,43^ Furthermore, huEGFRt expression can be monitored by fluorescence conjugated anti-huEGFR mAb and flow cytometry, giving it a secondary purpose as a trackable gene marker of vector modified cells.^42^

Other negative selection strategies of vector-modified HSPC have been developed using CD20 paired with rituximab, herpes simplex virus-thymidine kinase (HSV-TK) paired with ganciclovir, and inducible caspase 9 (iCas9) paired with AP1903 (Rimiducid) to induce dimerization.^59–61^ CTX-mediated negative selection of huEGFRt+ cells stands out as a promising safety switch for several compelling reasons. Unlike rituximab used to deplete CD20+ cells, CTX has not been shown to cause late onset neutropenia in clinical trials.^42,43,62,63^ CTX-mediated elimination of huEGFRt+ cells also holds several advantages to the HSV-TK system. The HSV-TK system paired with ganciclovir is only functional on proliferating cells, and a loss of sensitivity of ganciclovir could further stifle this method’s effectiveness.^64,65^ Studies have also indicated the potential for immunogenicity against HSV-TK, and its interference with host DNA repair, greatly enhancing the potential for unwanted cytotoxicity.^66,67^ The iCas9 system has shown promise, with exposure to AP1903 leading to elimination of 85-95% of circulating iCas9- transduced cells *in vitro* and *in vivo*.^68–70^ However, there are limitations to the iCas9 system’s practical application in clinical trials, as safety switches derived from non-human sequences will likely increase the risk of immunogenicity.^71–73^ The significant reduction of huEGFRt+ vector-modified cells by CTX in huBLT mice in this study combined with prior use in clinical studies suggests its efficacy as a safe and successful kill-switch system for HSPC-based gene therapies.

In summary, we provided a proof of concept that our newly developed multi-pronged anti-HIV-1 gene lentiviral vector with a safety kill switch could mediate efficient HSPC CD34+ transduction, engraftment, viral load reduction, and negative selection of vector modified cells *in vivo* in huBLT mice. Our studies performed in humanized BLT mice provide valuable insight on the development, protection, and efficacy of engineered T cells from anti-HIV-1 gene modified human HSPC developed through the human thymus. For clinical application, we recognize that there are still many obstacles to overcome. Further investigation of our strategies in more clinically relevant animal models, such as non-human primates, could provide us more clinically relevant results. We believe continuous improvements in the level of HIV-1 inhibition by enhancing the engraftment of anti-HIV-1 gene modified HSPC and ensuring safety will ultimately succeed us to translate our HSPC based anti-HIV-1 gene therapy into clinic.

## Materials and Methods

This study was carried out in strict accordance with the recommendations in the Guide for the Care and Use of Laboratory Animals of the National Institutes of Health (“The Guide”), and was approved by the Institutional Animal Care and Use Committees of the University of California, Los Angeles, protocol #ARC-2007-092. For humanized mice, all surgeries were performed under ketamine/xylazine and isoflurane anesthesia, and all efforts were made to minimize animal pain and discomfort.

### Vector Construction

Our vector construct backbone is derived from the “EQ” plasmid (generously provided by Satiro N. De Oliveira, UCLA, Los Angeles, California).^43^ We inserted our previously constructed CCR5sh1005, ^29^ D1D2CAR 4-1BB previously published by Zhen et al,^37^ and also the membrane anchored HIV-1 fusion inhibitor C46 previously published by Burke et al.^30^ Final optimized constructs also included the CD8 stalk element after the D1D2CAR 4-1BB extracellular domains and before the CD8 transmembrane domain in the D1D2CAR 4-1BB. Ubiquitin C or modified MLV long terminal repeat (MNDU3) promoters were used in each vector respectively to express D1D2CAR 4-1BB and huEGFRt. C46 was expressed using a truncated elongation factor-1a (EF1a) promoter. The final construct plasmids (M1 and U1) were purified using QIAquick Gel Extraction Kit (Qiagen, Hilden, Germany). Stellar competent cells from Takara Bio (Kusatsu, Shiga, Japan) were transformed with our constructed plasmids, and plasmid stocks were then produced using Macherey-Nagel Nucleobond Xtra Midi Kit (Macherey-Nagel, Düren, Germany).

### Cell culture

MT4-CCR5 cells are a human T-lymphotropic virus type 1-transformed human CD4^+^ T cell line that stably expresses CCR5, and were kindly provided by Dr. Koki Morizono (UCLA, Los Angeles). MT4-CCR5 cells were generated by transducing MT4 cells with a lentiviral vector expressing human CCR5 under the control of the internal SFFV promoter. These cells were cultured in RPMI-1640 (Life Technologies, Carlsbad, CA) supplemented with 10% fetal bovine serum (FBS), 2mM L-glutamine, 100 units/ml penicillin, and 100 μ Human primary peripheral blood mononuclear cells (PBMC) were isolated from whole blood from healthy donors obtained by the UCLA CFAR Virology Core Laboratory using Ficoll Paque Plus (GE Healthcare, Uppsala, Sweden). PBMC were cultured in RPMI-1640 supplemented with 20% FBS and GPS (20F RPMI). Primary human fetal liver derived (FL) CD34+ HSPC were isolated from human fetal liver obtained from the UCLA CFAR Gene and Cellular Therapy Core Laboratory (Los Angeles, California) or Advanced BioScience Resources, Inc (Alameda, CA) using a CD34^+^ microbead isolation kit (Miltenyi Biotec, Auburn, CA).

### Lentivirus Production

Lentiviral vectors were packaged with a VSV-G pseudotype using calcium phosphate transfection, then collected from transfected HEK293T-cells treated with chloroquine and concentrated in 1xHBSS via ultracentrifugation, as previously described.^74–76^ Titers for lentiviral vectors were determined using CTX-PE mAb staining and flow cytometry in vector-transduced HEK293T-cells and were based on huEGFRt expression at Day 3 post-transduction.

### *In-Vitro* HIV-1 inhibition assays

We established our vectors’ proof of concept to inhibit HIV-1 and target HIV-1 infection *in-vitro* in cell lines and primary cells prior to *in-vivo* experiments. For our HIV Inhibition assay, MT4-CCR5 cells were transduced with triple-anti-HIV vectors at MOI 1 and infected with R5- tropic HIV-1_NFNSX_ virus (MOI 1) or X4-tropic HIV-1_NL4-3_ virus (MOI 0.005) at 4 days post-transduction. Supernatants collected 7 days post-challenge were analyzed for p24 using Abcam HIV p24 ELISA kit (Abcam, Waltham, MA). To functionally qualify D1D2CAR 4-1BB, CD8+ T cells were isolated from healthy-PBMCs provided by UCLA CFAR virology core and expanded in RPMI 1640 (Gibco) with 10% fetal bovine serum (HyClone) containing IL7 (10ng/mL) and IL15 (5ng/mL). Cells were transduced with lentiviral vectors at MOI 3 and co-cultured with either unstimulated ACH2 cells or ACH2 cells stimulated overnight with PMA/Ionomycin (Invitrogen, Darmstadt, Germany) to increase HIV-1 gp120 surface protein.^77,78^ ACH2 cells are a cell line with a single integrated copy of HIV-1 strain LAI. Unstimulated and stimulated ACH2 cells were labeled with CellTrace™ Far Red (Invitrogen) before 16-hr co-culture followed by stain with Zombie (Aqua or Green) Fixable Viability (Biolegend, San Diego, CA) and KC57 antibody (Beckman Coulter, Indianapolis, IN) to detect Gag+ ACH2 cells.

Specific killing was calculated as follows: % specific killing =(%live Gag+ ACH2 cells co-cultured with untransduced cell—%live Gag+ ACH2 cells co-cultured with vector-transduced cells)/ %live Gag+ ACH2 co-cultured with untransduced cell × % Gag+ in ACH2 cells alone.

### Lentiviral Vector Transduction of HSPC for *in vivo* experiments

Fetal liver-derived CD34+ HSPCs were resuspended in Yssel’s medium (Gemini Bio Products) with 2% BSA (Sigma-Aldrich) and seeded into 20 µg/mL RetroNectin (Clontech Laboratories)-coated plates. After 1 hour of incubation at 37LJ lentiviral vectors at MOI 3 and cultured overnight at 37LJ CD34+ HSPCs were transplanted into NSG mice. An aliquot of the transduced CD34+ HSPCs were cultured in 10F RPMI, supplemented with cytokine stimulations (SCF, Flt-3, TPO; PeproTech) at a concentration of 50 ng/mL for 3 days. The efficiencies of vector transduction were evaluated by flow cytometry (Fortessa flow cytometers, BD Biosciences) and/or by vector copy number (VCN) using digital PCR as described below (ThermoFisher QuantStudio 3D Digital System/Quantstudio Absolute Q Digital PCR system).

### Humanized BLT Mouse Construction

NSG (NOD/SCID/IL2rγ -/-) mice were used to generate humanized BLT mice and housed according to UCLA Humanized Mouse Core Laboratory procedures as previously described. ^31^ Human fetal thymus and fetal liver were obtained from Advanced Bioscience Resources (ABR). Fetal tissues were obtained without patient identifying information. Written informed consent was obtained from patients for the use of tissues for research purposes. Briefly, one day before transplant, CD34+ cells were isolated from fetal livers using anti-CD34+ magnetic bead-conjugated monoclonal antibodies (Miltenyi Biotec) and transduced with vectors described above. NSG mice were conditioned with sub-lethal body irradiation (270 cGy Cesium- 137). On the day of transplant, an equal mixture of non-transduced or vector-transduced FL- CD34+ cells (∼ 0.5x10^6^ per mouse) and CD34- cells (∼4.5 x10^6^ per mouse) were mixed with 5μL of Matrigel (BD Biosciences) and implanted with a piece of thymus under the kidney capsule. Mice were then injected with non-transduced or vector-transduced CD34+ HSPCs (∼ 0.5x10^6^ per mouse) using a 27-gauge needle through the retro-orbital vein plexus. At 8–10 weeks post-transplantation, blood was obtained from each mouse by retro-orbital sampling and peripheral blood mononuclear cells were analyzed by flow cytometry to quantify human immune cell engraftment.

### Colony-forming unit (CFU) assays

Colony-forming units were assayed by culturing transduced and non-transduced FL- CD34+ cells 3 days after transduction in triplicate in a 6 well plate (ThermoFisher) using complete methylcellulose (MethoCult H4435 Enriched, Stem Cell Technologies). 14 days later, colony-forming units (CFU) in each well were then counted by light microscopy, and the colony type was scored based on morphology. Proportions of differentiated hematopoietic colonies = 100% × (each colony type CFU counted/total CFU counted) and calculated from each well from triplicates.^79^ Total CFU counts ranged from 30-75 in each well.

### HIV-1 Infection and viral load analysis

NSG huBLT mice were injected with R5 tropic HIV-1_NFNSX-SL9_ (MOI 5) (200 ng p24) retro-orbitally 11 weeks post-vector-modified HSPC transplant ^74^. Mice were bled retro-orbitally every 2 weeks after infection, and blood samples were analyzed for HIV-1 viral load via RT-PCR. HPSC engraftment was assessed by vector copy number assay via digital PCR, and cell lineage differentiation and transgene expression were measured via flow cytometry.

### Depletion of huEGFRt+ transduced cells via Cetuximab

At week 13-14 post-transplantation of vector-modified HSPC, mice were separated into two groups with one group to receive CTX treatment (CTX+) alongside human NK cells and huIL-15 treatment (n=5) and the other group to be left untreated (n=4). Vector-modified huEGFRt+ HSPC transplanted huBLT mice were treated with cetuximab (Erbirtux^TM^) at a concentration of 1mg per mouse intraperitoneally for 11 consecutive days. Because of the lack of efficient development of human NK cells in NSG mice, which was hypothesized to be the result of a lack of IL15,^80^ and to facilitate antibody dependent cellular toxicity, we injected a dose of 5×10^6^ human NK cells isolated from healthy PBMCs one day before the first CTX treatment and a second dose of 5×10^6^ human NK cells from the same donor on day 7 of CTX treatment. Lentiviral vectors expressing IL-15 (2.5×10^5^ IU/mouse) were injected retro-orbitally on day 7 of CTX treatment. huEGFRt expression and absolute cell count were monitored in multi-human cell lineages by staining peripheral blood cells with CD45-, CD3-, CD19-, CD4-, and CD8- specific monoclonal antibodies of peripheral blood prior to flow cytometry analysis at 3 weeks before CTX treatment and 1 and 4 weeks post-CTX treatment. We developed this strategy to augment negative selection results in animals lacking circulating NK cells and supportive IL-15, as NK cells serve a critical role in CTX-mediated ADCC of huEGFRt+ cells (**Figure S7**).

### Analysis of tissue from transduced huBLT mice

Humanized BLT mice were sacrificed at week 18-19 post-transplant, and the spleen and BM tissues were harvested. Tissue samples were collected in MACS tissue storage solution (Miltenyi Biotec, 130-100-008) at necropsy and processed immediately for single cell isolation as described previously.^31,37^ Isolated cells were stained for surface markers and analyzed by flow cytometry or vector copy number was determined by digital PCR.

Single-cell suspensions prepared from peripheral blood, spleen, or BM of huBLT mice were stained for surface markers and acquired on a LSRFortessa flow cytometer (BD Biosciences). The following antibodies were used: CD45-eFluor 450 (HI30, eBioscience), CD3- APC H7 (SK7, Pharmingen), CD4-APC (OKT4, eBioscience), CD8-PerCP Cy5.5 (SK1, BioLegend), CD19-Brilliant Violet 605 (HIB19, BD Horizon), EGFR-PE (Hu1, R&D Systems), and Countbright beads (Invitrogen). Red blood cells were lysed with RBC Lysis Buffer (Biolegend) after cell surface marker staining. Stained cells were fixed with 2% formaldehyde in PBS. The data were analyzed by FlowJo v.10 (Tree Star) software.

### Determination of vector copy number (VCN)

Cell pellets from 25 μL peripheral blood, spleen, or BM of huBLT mice were lysed with 5μL of 0.2M NaOH in a 75□ water bath for 5 min. Cell lysates were cooled in a 4°C refrigerator for 5 min 45μL Tri-HCl was added to neutralize the lysates. The lysate cells were directly used in dPCR set at 96°C for 10 min, followed by 42 cycles of denaturation at 98°C for 15 s, annealing at 60°C for 2 min, and a final extension at 60°C for 2 min. The primers and probe specific to WPRE were customized by ThermoFisher, which are primer sequence 1, 5’ - CCTTTCCGGGACTTTCGCTTT-3’, primer sequence 2, 5’-GCAGGCGGCGATGAGT-3’, and probe 5’-(FAM)-CCCCCTCCCTATTGCC-3’. The primers and probe specific to β purchased from ThermoFisher (cat#4448489). Average VCN was determined by multiplex dPCR of the WPRE sequence in the vector and normalized to the cell housekeeper gene β globin.

### Statistical Analysis

Statistical analysis was performed using software Prism. Mann-Whitney U test was used for nonparametric testing of independent groups, and student’s t-test were used for parametric testing of independent groups. Statistical significance was evaluated as *p <0.05. Other significance levels are indicated as follows: **p <0.01, ***p <0.001, and ****p <0.0001.

## Data Availability Statement

The data supporting the findings of this study are available from the corresponding author upon reasonable request.

## Author contributions

Authors QG, KP, and JZ contributed equally to this manuscript and are all considered co-first authors. Authors AB, NJ, and GC contributed to experimental data collection and manuscript revision. Authors AZ and DSA gave invaluable guidance and feedback throughout the preparation of this manuscript. All authors had the opportunity to review the manuscript prior to submission.

## Acknowledgements

We would like to thank Sarah Schroeder, Christina Zakarian, Martin Zakarian, Kory Hamane, and Dr. Scott Kitchen, Valerie Rezek and staff at UCLA Humanized Mouse & Gene Therapy Core for their technical support. We would also like to thank Dr. Chris Peterson, Dr. Paul Krogstad, and Dr. Irvin Chen for their feedback and revisions on drafted versions of manuscript. This research was supported by the NIAID 1U19 AI149504, the UCLA-CDU Center for AIDS Research NIH/NIAID AI152501, the NIAID R01AI172727 to AZ, the NIDA R01DA-52841 to AZ, the amfAR 110304-71-RKRL and 110395-72-RPRL to AZ, the James B. Pendleton Charitable Trust, and UCLA AIDS institute. The vector maps and experimental design figures were created with biorender.com.

