## Supplemental Information for "Anti-HIV-1 HSPC-based gene therapy with safety kill switch to defend against and attack HIV-1 infection"

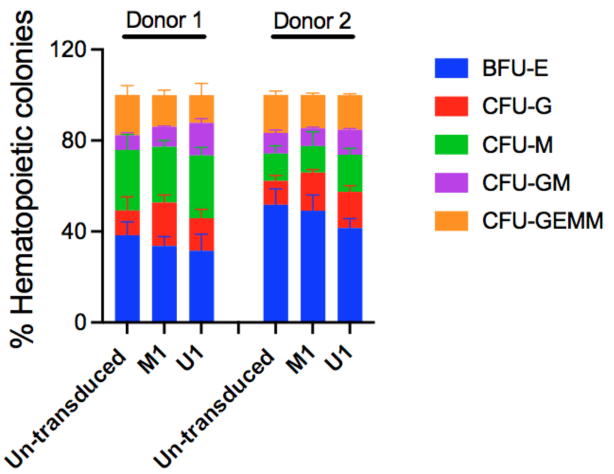


**Figure S1.** Colony-forming units (CFUs) of untransduced, M1, and U1 vector-transduced FL-CD34+ cells after 14 days of culture. % of burst-forming unit-erythroid (BFU-E), colony-forming unit-granulocytes/macrophages (CFU-G, CFU-M, CFU-GM), and multipotential progenitor cells (CFU-GEMM) = 100 × (each colony type CFU counted/total CFU counted). Data were collected from triplicates from two independent donors.

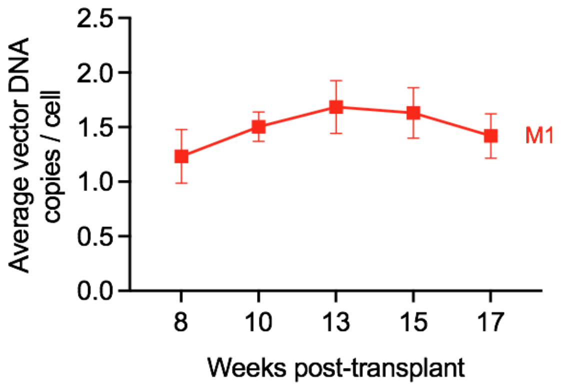


**Figure S2.** Vector copy number data in replicate (donor 2) huBLT mouse experiment with M1 vector-transduced HSPC transplanted huBLT mice. Vector-marking levels were determined in peripheral blood cells from 8 weeks to 17 weeks post-transplant by digital PCR. Average vector DNA copies were calculated by VCN = [WPRE DNA copies in vector DNA/ul]/[human β-globin copies/ul/2]. Dots and error bars show Mean ± S.E.M, respectively.

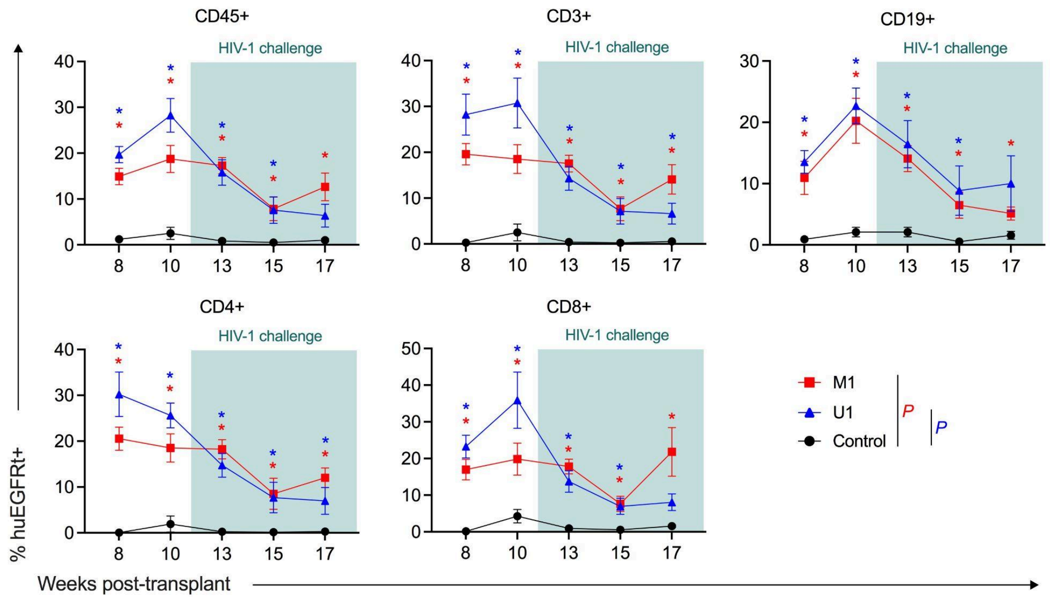


**Figure S3.** Percentage of huEGFRt expressing cells (huEGFRt+) in human hematopoietic lineages of peripheral blood in HSPC transplanted huBLT mice. Dots and error bars show Mean ± S.E.M, respectively. Mann-Whitney U test was performed to calculate significance, *p <0.05.

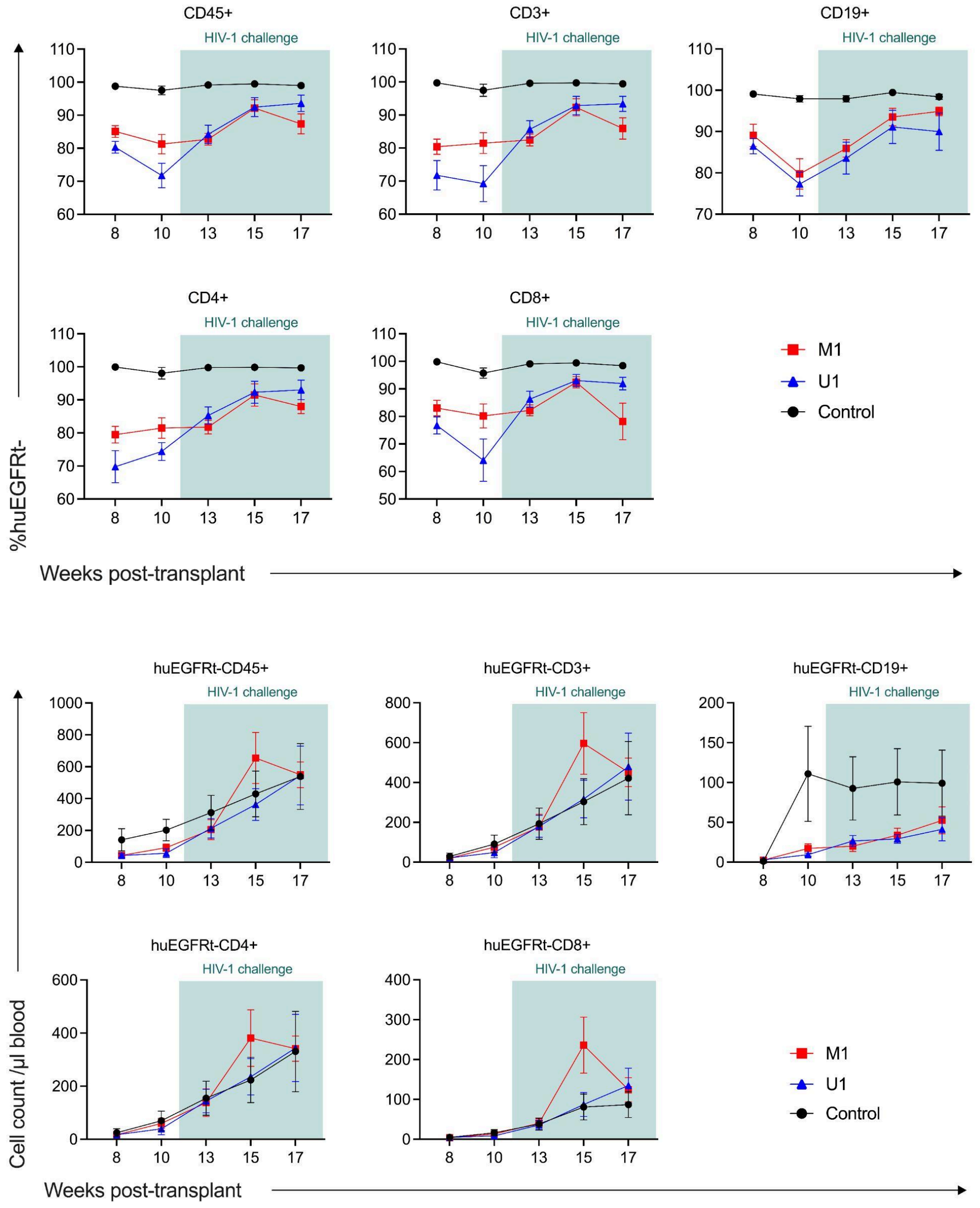


**Figure S4.** Percentage and absolute cell count/µL of huEGFRt non-expressing cells (huEGFRt-) in human hematopoietic lineages of peripheral blood in HSPC transplanted huBLT mice. Dots and error show Mean ± S.E.M, respectively.

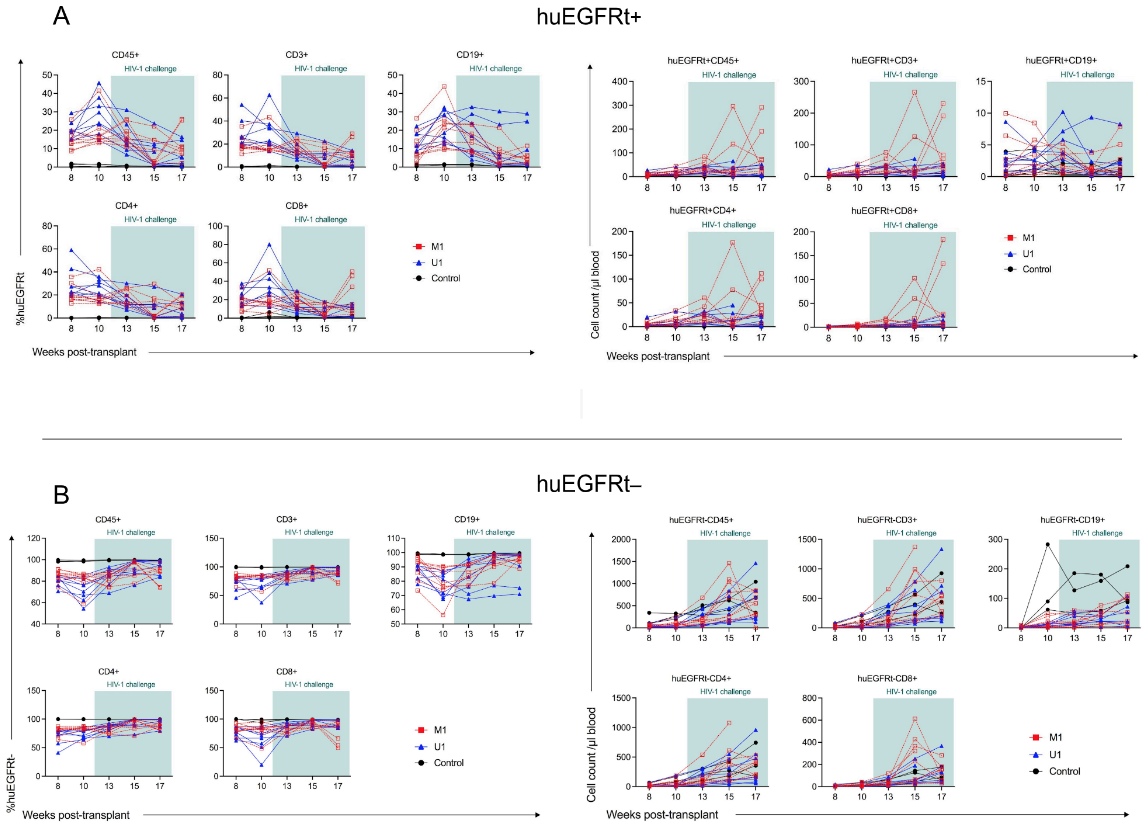


**Figure S5.** (A) Longitudinal gene marking (huEGFRt+) or (B) huEGFRt- marking in percentages and cell count/μL of peripheral blood in human multilineage hematopoietic populations from individual mice. Each dot represents an individual mouse from the corresponding group.

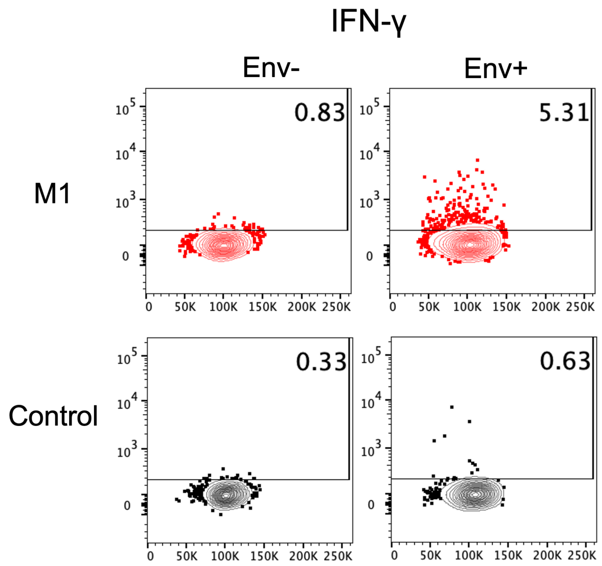


**Figure S6.** Representative *ex vivo* cytokine production from vector-transduced CD8+ T splenocytes. Effector cells from vector transduced HPSC transplanted huBLT mice were co-cultured with Env+ target cells (PMA/ionomycin activated ACH2 cells) or unstimulated Env- cells (medium only) as a negative control *ex vivo*. Data was collected from our replicate huBLT mice experiment (donor 2). Cells were collected at time of mouse necropsy (week 20 post-transplant) and cytokine production was measured by flow cytometry.

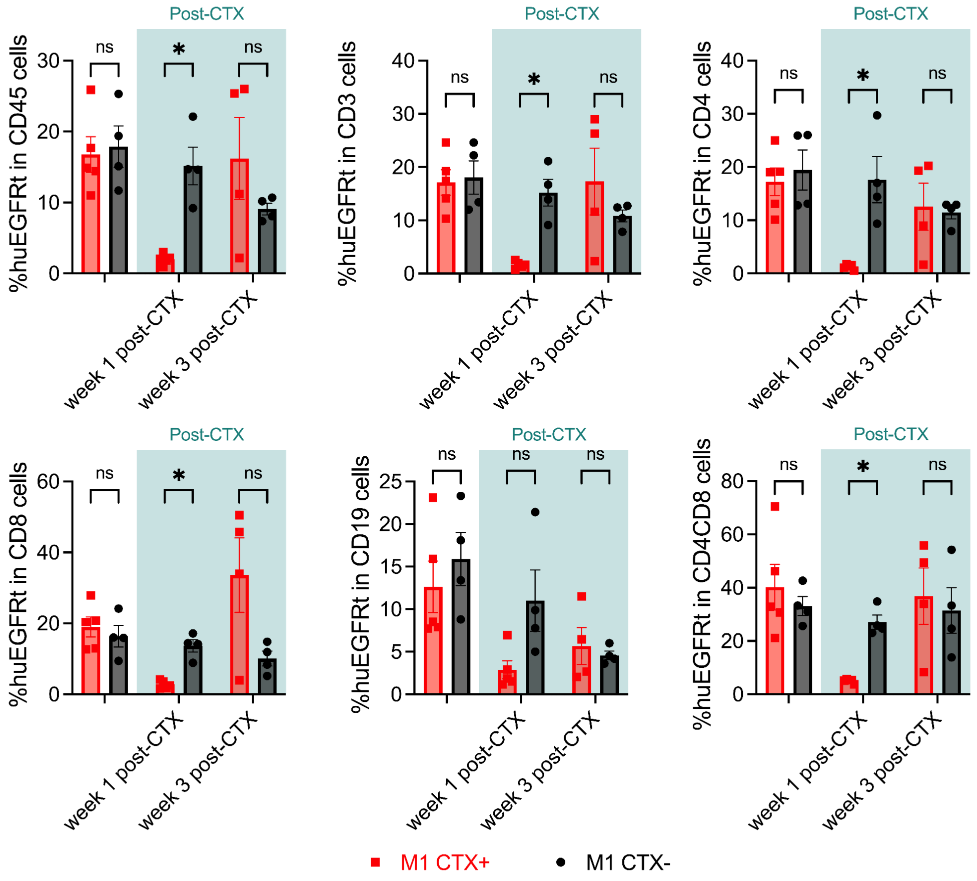


**Figure S7.** HuEGFRt expression level in multilineage human lymphocytes (CD45), T-cells (CD3, CD4, and CD8) and B-cells (CD19) in CTX treated and untreated huBLT mice that were not infused with human NK cells. Blood was collected 1 week before CTX treatment, and 1 and 3 weeks post-onset of CTX treatment. HuEGFRt expression was measured by flow cytometry using mAb CTX-PE. Dots and error bars show Mean ± S.E.M, respectively. Mann-Whitney U test was performed to calculate significance. NS = not significant, and *p <0.05.

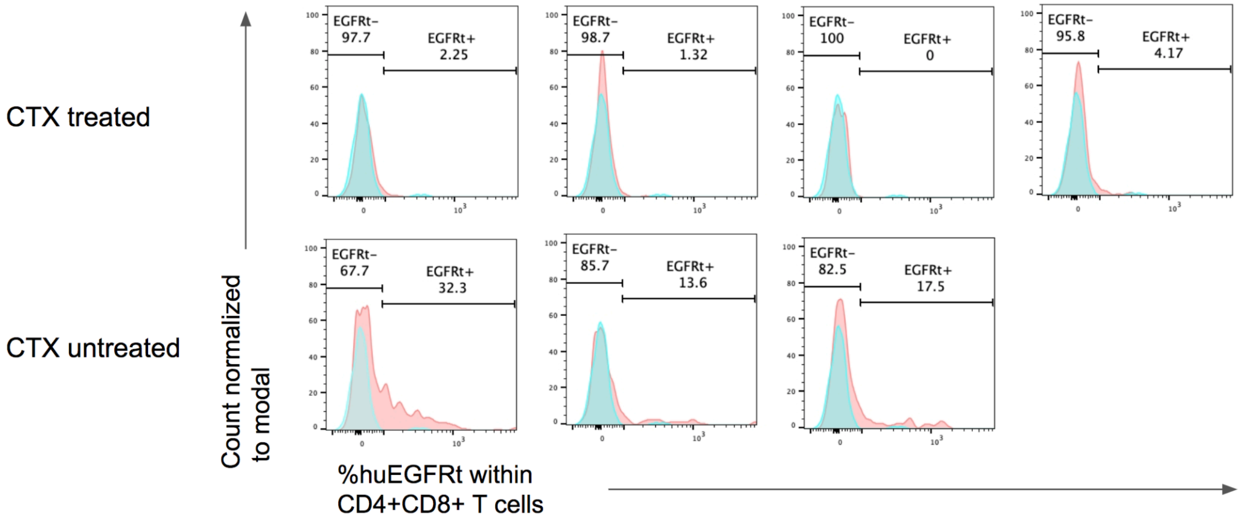


**Figure S8.** Percentage expression of huEGFRt+ vs huEGFRt- cells within CD4+CD8+ T cell population of splenocytes collected from CTX treated and untreated M1 huBLT mice (as shown in Figure 3D). Each plot represents an individual mouse of the treatment group. Blue plot represents a single untransduced huBLT mouse used as a negative control.

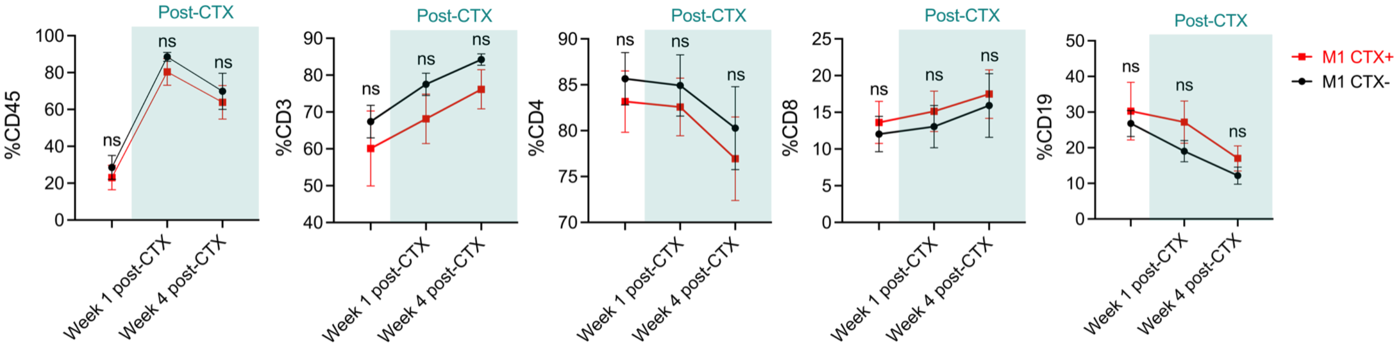


**Figure S9.** Human multilineage hematopoietic cell reconstitution in peripheral blood of CTX treated and untreated huBLT mice with huEGFRt+ cell lineages. Blood samples were collected at 3 weeks before CTX treatment, and 1 and 4 weeks post-treatment. Surface markers of human lymphocytes (CD45), T-cells (CD3, CD4, and CD8) and B-cells (CD19) were stained and measured by flow cytometry.

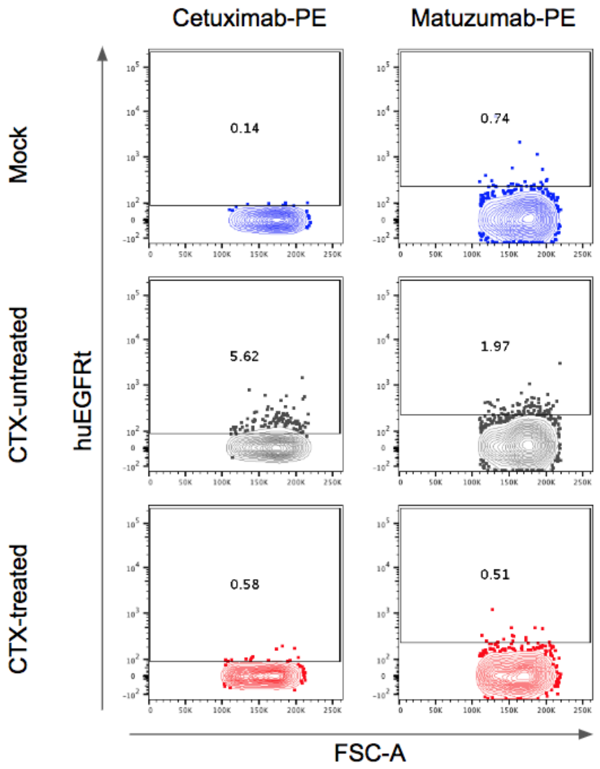


**Figure S10.** Representative flow cytometry data showing huEGFRt expression stained in splenocytes. “CTX-untreated” splenocytes were transduced with vector M1 but not treated with CTX, while “CTX-treated” splenocytes were transduced with vector M1 and treated with CTX. “Mock” splenocytes were not transduced with any vector and were not treated with CTX, serving as a negative control. Splenocytes were stained with two different mAbs, CTX-PE and non-competing anti-huEGFR mAb MTZ-PE. MTZ-PE was utilized to confirm the depletion of huEGFRt expressing population. Expression was measured by flow cytometry.

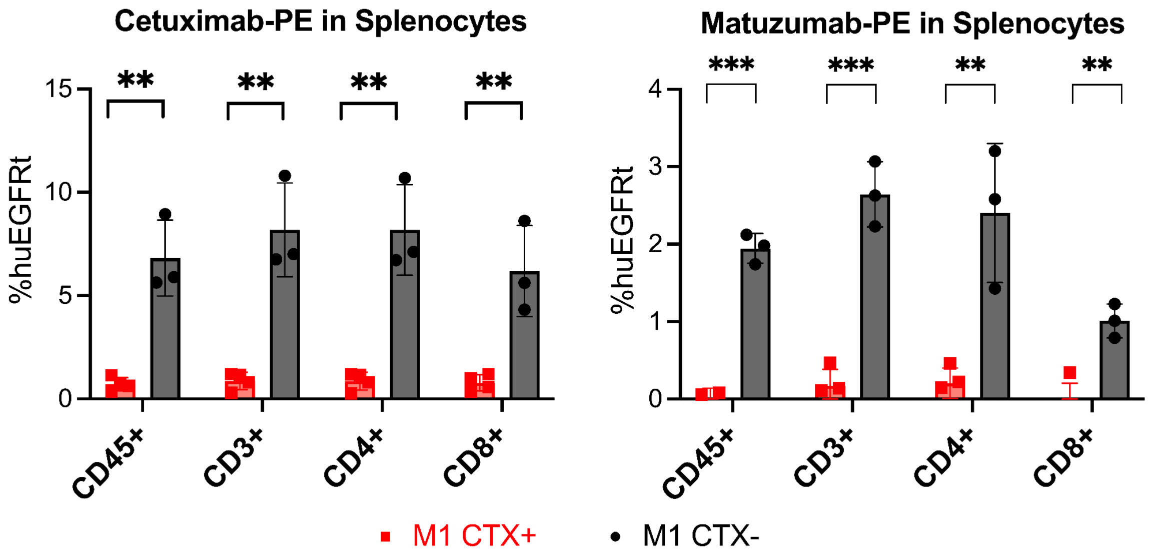


**Figure S11.** HuEGFRt expression level across multilineage human cell populations including human lymphocytes (CD45), T-cells (CD3, CD4, and CD8) and B-cells (CD19) in splenocytes collected from CTX-treated and untreated huBLT mice. Samples were stained with two different mAbs CTX-PE (as shown in Figure 3C) and non-competing anti-huEGFR mAb MTZ-PE. MTZ-PE was utilized to confirm depletion of huEGFRt expressing population, and measured by flow cytometry. Dots and error bars show Mean ± S.E.M, respectively. Student's t-test was performed to calculate significance. **$p < 0.01$, and ***$p < 0.001$.
